# BCR ligation selectively inhibits IgE class switch recombination

**DOI:** 10.1101/2024.09.18.613749

**Authors:** Adam K. Wade-Vallance, Zhiyong Yang, Jeremy B. Libang, Ananya R. Krishnapura, James B. Jung, Emily W. Matcham, Marcus J. Robinson, Christopher D. C. Allen

## Abstract

Mechanisms that restrict class switch recombination (CSR) to IgE limit the subsequent production of IgE antibodies and therefore the development of allergic disease. Mice with impaired B cell receptor (BCR) signaling have significantly increased IgE responses, consistent with a role for BCR signaling in IgE regulation. While prior work focused on BCR signaling in IgE-expressing cells to explain these findings, it has been reported that BCR signaling can reduce CSR. Therefore, we investigated the possibility that IgE CSR might be particularly sensitive to inhibition by BCR signaling in unswitched B cells. We found that immunization of mice with high-affinity antigen resulted in reduced representation of IgE-expressing cells among germinal center B cells and plasma cells relative to a low-affinity antigen. Mechanistic experiments with cultured mouse B cells demonstrated that BCR ligands selectively inhibited IgE CSR in a dose-, affinity-, and avidity-dependent manner. Signaling via Syk was required for the inhibition of IgE CSR following BCR stimulation, whereas inhibition of the PI3K subunit p110δ increased IgE CSR independently of BCR ligation. The inhibition of IgE CSR by BCR ligands synergized with IL-21 or TGFβ1. BCR ligation also inhibited CSR to IgE in human tonsillar B cells, and this inhibition was also synergistic with IL-21. These findings establish that IgE CSR is uniquely susceptible to inhibition by BCR signaling in mouse and human B cells, with important implications for the regulation and pathogenesis of allergic disease.

## Introduction

Type-1 hypersensitivity responses in allergic disease result from the production of IgE antibodies specific for food and environmental antigens. Prior to IgE production, B cells must first undergo IgE class switch recombination (CSR). Whereas IgE CSR can be readily induced by stimulating B cells in cell culture, the abundance of IgE in serum is orders of magnitude less than that of IgG. The scarcity of IgE *in vivo* implies the existence of regulatory mechanisms absent from typical *in vitro* systems.^1^ For example, our prior work identified IL-21 as a major negative regulator of IgE CSR.^2^ An additional potential source of regulation is cognate antigen. Indeed, when cognate antigen directly ligates B cell receptors (BCRs) on IgE plasma cells (PCs; used to collectively refer to both plasma cells and plasmablasts), this results in their elimination.^3^ Stimulation of the BCR can also broadly inhibit class-switching,^4–7^ and prior studies from our lab and others found selectively increased IgE responses in mice with impaired BCR signaling.^8–10^ However, evidence for whether BCR stimulation might selectively affect IgE CSR is mixed.^6,11^

B cell antigen encounter initiates a complex array of BCR signaling pathways that coordinate a variety of cellular responses. The incipient events of BCR signaling include the formation of the “BCR signalosome” through a series of interactions between Syk, BLNK, Btk, and PLCγ2, among others.^12^ This process is reinforced through the activation of PI3K (composed of p110 catalytic and p85 regulatory subunits) to produce phospholipids that recruit several BCR signalosome components and other molecules to the plasma membrane.^12^ Mice with disrupted p110δ (the most abundant p110 isoform in B cells)^13–15^ signaling produce exaggerated IgE responses to type-2 immunizations.^8,10,16^ These observations may relate to effects on IgE CSR, as CSR is broadly inhibited in mice with enhanced PI3K activity, and treating mice or cells with PI3K inhibitors increases IgE.^4,16–20^ Increased IgE responses have also been reported in mice with heterozygous mutations in *Syk*, homozygous mutations in *Blnk*, or upon pharmacologic inhibition of Btk.^10^ Syk, BLNK, and Btk are known to be important in cells that have already undergone IgE CSR due to their roles in antigen-independent signaling of the IgE BCR^8,10^ and in BCR ligation-induced IgE PC apoptosis.^3^ It is unclear if these molecules also play a role in the inhibition of IgE CSR downstream of BCR stimulation.

Here, we investigated the role of BCR signaling in IgE CSR. We first used genetic and comparative immunization-based strategies in mice to demonstrate that stronger BCR:antigen interactions resulted in reduced representation of IgE cells in the GC and among extrafollicular PCs. BCR stimulation inhibited IgE CSR to a greater extent than IgG1 CSR, and CSR inhibition was dose-, affinity-, and avidity-dependent. We identified a selective effect of BCR stimulation on ε germline transcripts, implicating a transcriptional mechanism for IgE CSR inhibition. At the level of signaling molecules, Syk was required, and PKC signaling was sufficient, for the selective inhibition of IgE CSR following BCR stimulation. Interestingly, p110δ was not required and instead suppressed IgE CSR independently of BCR stimulation. We found synergistic inhibition of IgE CSR with BCR stimulation and either IL-21 or TGFβ1. BCR stimulation also selectively inhibited IgE CSR in human B cells, which was also synergistic with IL-21.

## Results

### BCR stimulation strength inversely correlates with IgE-switched cells

To query how BCR:antigen binding strength might selectively regulate IgE CSR *in vivo*, we performed adoptive transfers of Hy10 B cells whose BCRs are specific for avian egg lysozyme. Congenic recipient mice were then immunized with antigen of low or high affinity for the Hy10 BCR (duck egg lysozyme [DEL] or hen egg lysozyme [HEL], respectively).^21^ One week later, IgE, but not IgG1, cells were less frequent among both GC B cells and PCs in the draining lymph nodes (dLNs) of recipients immunized with HEL compared to DEL (Figure 1A). These data provide evidence for a selective. inhibitory effect of antigen affinity on the frequency of IgE-expressing cells *in vivo*.

**Figure 1.**
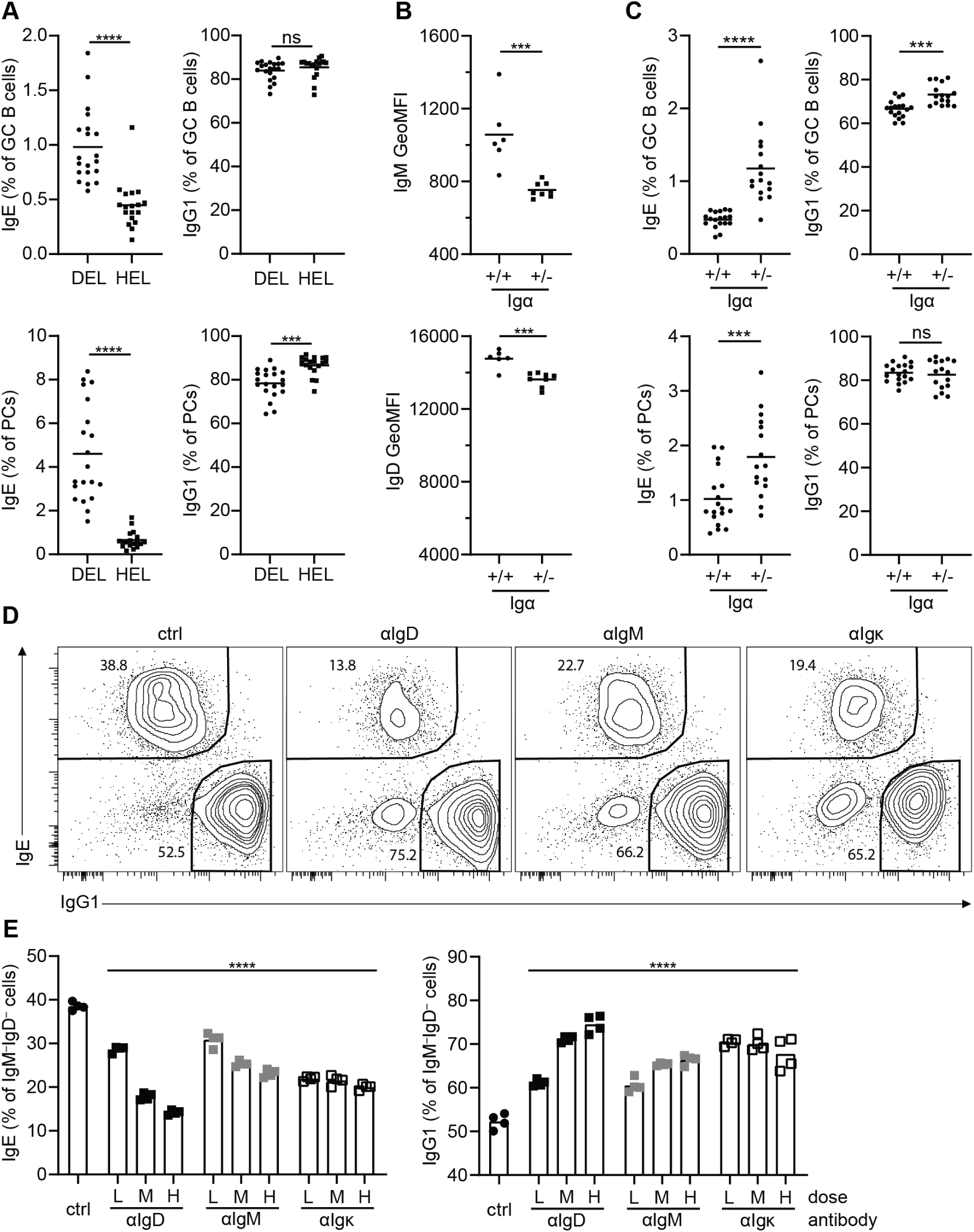
BCR stimulation reduces the representation of IgE cells *in vivo* and *in vitro*. **(A)** Hy10 B cells were adoptively transferred into congenically-marked recipient mice that were then immunized with ovalbumin-conjugated DEL or HEL in alum adjuvant. Shown are the proportions of IgE cells (left column) and IgG1 cells (right column) within the GC (top row) and PC (bottom row) compartments of transferred cells in the dLN at d7, quantified by flow cytometry. (B) Quantification of surface IgM (top) and IgD (bottom) levels on follicular B cells from wildtype or Igα^+/-^ mice by flow cytometry. (C) Wildtype and Igα^+/-^ mice were immunized subcutaneously with NP-CGG in alum adjuvant and the resultant immune response in the dLN at d7 was analyzed by flow cytometry. Plots are laid out as described for panel A. (D-E) Wildtype mouse B cells were cultured with IL-4, αCD40, and the indicated treatments for 4 days prior to analysis by flow cytometry. (D) Representative flow cytometry plots of IgE and IgG1 staining among class-switched (IgM^−^IgD^−^) cells according to treatment condition (from left to right; ctrl [GGG], αIgD, αIgM, αIgκ; all at 1 μg/mL). (E) Quantification of the effects of treatment with low (L; 100 ng/mL), medium (M; 300 ng/mL), or high (H; 1 μg/mL) doses of αIgD (black squares), αIgM (grey squares), αIgκ (white squares), or ctrl (GGG; 1 μg/mL; black circles) antibodies on the proportions of IgE (left) and IgG1 (right) cells among class-switched (IgM^−^IgD^−^) cells. (A-C, E) Dots represent samples from individual mice and bars represent the mean values. ns, not significant; *, P < 0.05; **, P < 0.01; ***, P < 0.001; ****, P < 0.0001 (unpaired t test [A-C], one-way repeated measures ANOVA [E] with Dunnett’s post-test comparing each condition to the control with the Holm-Sidak correction for multiple comparisons). Results are pooled from three (A) or two (B, C, E) independent experiments or are representative of two independent experiments (D).

We next sought to determine if modulating BCR signaling strength through the surface expression level of the BCR would impact IgE switching. Igα (aka CD79a / Mb1) is one of two obligate ITAM-containing protein components of the BCR, and is also critical for BCR surface expression.^10,22^ We therefore reasoned that Igα-heterozygous mice might have reduced surface BCR expression. Indeed, Igα^+/-^ naïve follicular B cells had reduced surface IgM and IgD relative to WT cells (Figure 1B). Following immunization, Igα^+/-^ mice had increased frequencies of IgE cells within both the GC and PC compartments (Figure 1C). Interestingly, these mice also had a subtle increase in IgG1 GC B cells and decrease in IgG1 PCs, consistent with previously described roles for BCR signaling in PC differentiation.^23,24^ Overall, these data expand upon prior findings that weakened BCR signaling results in greater IgE responses^8,10^ and are consistent with the hypothesis that IgE CSR is especially susceptible to inhibition by BCR ligation.

To dissect the effect of BCR ligation on IgE CSR we induced B cells, purified from murine splenocytes, to undergo CSR in cell culture using IL-4 and αCD40. BCR stimulation with an anti-BCR antibody (goat anti-mouse IgD [αIgD]), but not control antibody (goat gamma globulin [GGG]), resulted in a greatly reduced frequency of IgE cells in the culture (∼4.3-fold), with a lesser reduction in IgG1 cells (∼1.2-fold, Figure S1A-B). As we observed an overall reduction in class-switching after BCR stimulation (Figure S1C-D), we examined the frequency of IgE cells among class-switched cells and found that BCR stimulation resulted in greatly reduced representation of IgE cells within the class-switched compartment, while the representation of IgG1 cells was increased (Figure S1E-F). These data suggest that BCR stimulation has a selective negative impact on IgE CSR.

We next asked whether these findings were generalizable to different anti-BCR antibodies as well as cognate antigen. While αIgD treatment resulted in the greatest dose-dependent reduction in IgE of anti-BCR antibodies we measured, both goat anti-mouse IgM (αIgM) and goat anti-mouse Ig kappa light chain (αIgκ) produced significant reductions in IgE (Figure 1D-E). To investigate the impact of cognate antigen on IgE CSR, we purified and cultured Igλ light chain-expressing cells from B1-8i mice.^25^ B1-8i mice have a pre-arranged heavy chain VDJ that endows most B cells expressing λ light chains with binding specificity towards the hapten 4-hydroxy-3-nitrophenyl (NP) and with 20-fold higher binding strength towards the hapten 4-hydroxy-3-iodo-5-nitrophenyl (NIP).^26^ We assessed the impact of cognate antigen dose, affinity, and avidity on IgE and IgG1 CSR by treatment with different doses of high-valency, higher-affinity antigen (NIP_24_BSA); high-valency, lower-affinity antigen (NP_25_BSA); or low-valency, lower-affinity antigen (NP_4_BSA). As we previously found that cognate antigen could eliminate IgE PCs,^3^ we focused our present investigation on B cells by including antigen from the beginning of cell culture, using substantially lower doses, and performing our analysis prior to the bulk of PC differentiation. We found that all three cognate antigens resulted in dose-dependent reductions in the representation of IgE-expressing cells (Figure 2A), although the potency of the different cognate antigens varied over orders of magnitude according to their affinity and valency. Notably, the representation of IgE cells in culture was more sensitive to inhibition by cognate antigen than IgG1, as revealed by intermediate doses of all three cognate antigens which reduced the representation of IgE but not IgG1 cells. Examining the representation of IgE- and IgG1-switched cells among class-switched (IgM^−^IgD^−^) cells revealed that, even with powerful BCR ligation, the representation of IgE cells was reduced more greatly relative to other class-switched cells, whereas the representation of IgG1 cells was maintained or increased (Figure 2B). To quantitate the relative effects of cognate antigen on IgE and IgG1 CSR, we performed a normalized analysis (Figure 2C). This analysis confirmed that for each cognate antigen there were intermediate doses which led to decreases in IgE cells, but not IgG1 cells. While we observed reductions in IgG1 cells at high doses of antigen, these doses also resulted in greater reductions in IgE cells. Overall, these data support that ligating BCRs on B cells preferentially reduces IgE CSR.

**Figure 2.**
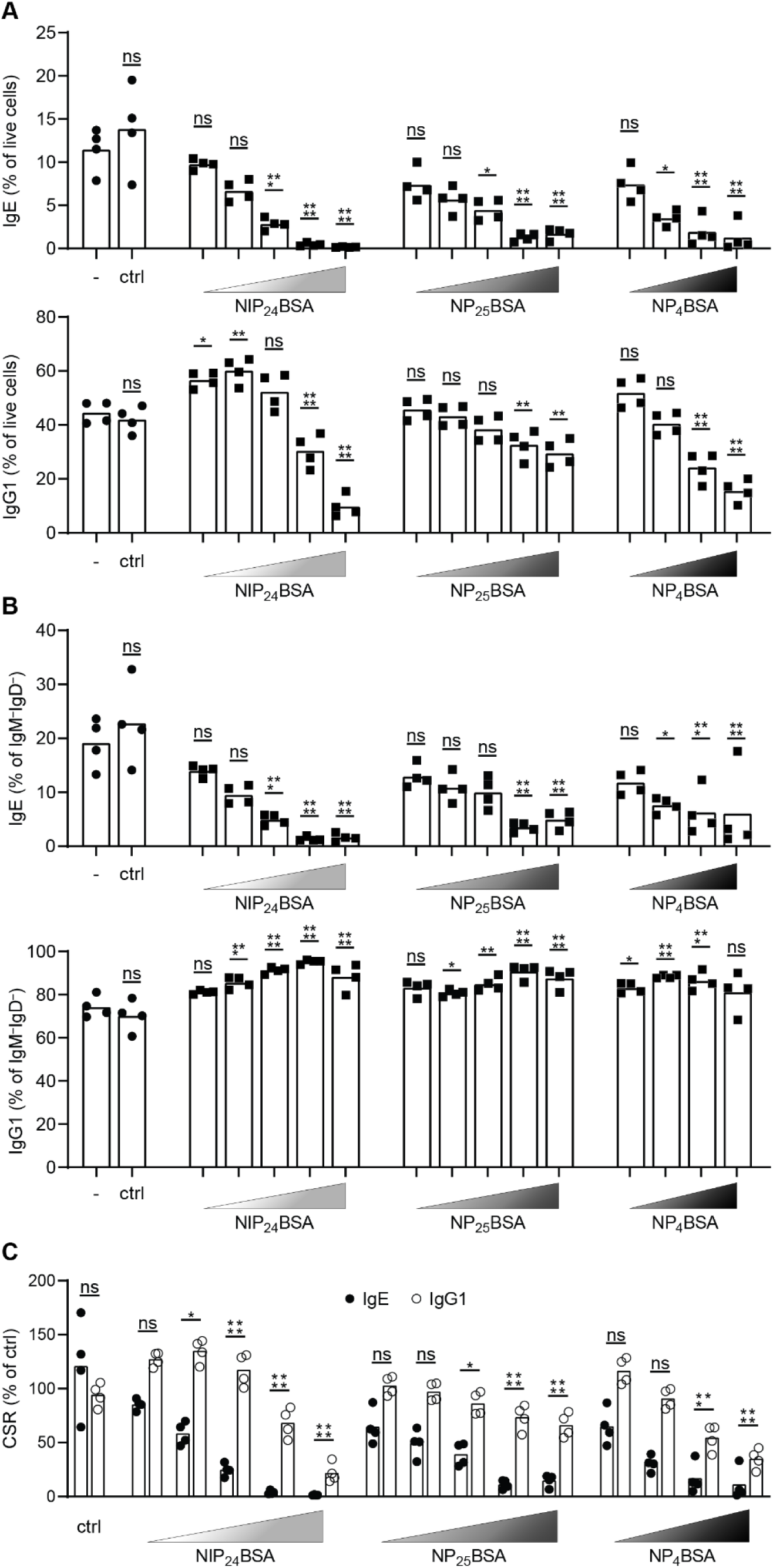
B cell culture with cognate antigen reduces yields of IgE cells. (A-C) Purified B1-8i B cells were cultured with IL-4 and αCD40 for 4 days with control (‘-’ is no treatment, ‘ctrl’ is BSA at 50 ng/mL; black circles) or cognate antigen (NIP_24_BSA, NP_25_BSA, or NP_4_BSA; black squares) prior to analysis by flow cytometry. Doses of cognate antigen ascend from left to right as represented by the gradient triangles, exact doses are as follows (ng/mL): NIP_24_BSA; 0.002, 0.005, 0.015, 0.05, 0.25 | NP_25_BSA; 0.016, 0.08, 0.4, 2, 10 | NP_4_BSA; 20, 200, 1.25 × 10^3^, 1 × 10^4^. (A-B) Quantification of the proportion of IgE (top) or IgG1 (bottom) cells among all live cells (A) or within the class-switched (IgM^−^IgD^−^) compartment (B) as assessed by flow cytometry. (C) Quantification of the frequency of IgE cells (black circles) and IgG1 cells (white circles) among live cells in the antigen-treated condition as a fraction of their frequency among live cells in the control condition. (A-C) Dots represent samples from individual mice and bars represent the mean values. ns, not significant; *, P < 0.05; **, P < 0.01; ***, P < 0.001; ****, P < 0.0001 (one-way repeated measures ANOVA with Dunnett’s post-test comparing each condition to the untreated control with the Holm-Sidak correction for multiple comparisons). Results are representative of two independent experiments.

### BCR stimulation inhibits IgE CSR by downmodulating ε germline transcription

We considered multiple possibilities by which stimulating the BCR could lead to reduced numbers of IgE cells. As class-switching is linked to cell division, and IgE requires more cell divisions to emerge relative to IgG1,^27^ one possibility is that BCR signaling limits the opportunity for IgE CSR to occur by reducing proliferation. To distinguish potential effects on proliferation versus CSR we loaded B cells with CellTrace Violet (CTV) and examined at each division number whether BCR stimulation affected the fraction of IgE or IgG1 cells. CTV fluorescence is reduced 2-fold with each cell division, and after four days of culture, we were able to visualize seven distinct populations of B cells based on the intensity of their CTV staining (Figure 3A). Whereas IgG1 cells were first detectable after 3 divisions, IgE cells remained absent until 5 divisions, consistent with prior work (Figure 3B-C).^27^ Relative to control treatment, BCR stimulation resulted in a reduction in the fraction of cells switched to IgE across all cell divisions at which they were present (Figure 3B). This finding suggests that BCR stimulation reduces IgE CSR directly rather than as a secondary outcome of impaired proliferation. Meanwhile, BCR stimulation resulted in a mild reduction in the fraction of IgG1-switched cells at divisions 2 through 5, but IgG1 CSR normalized and trended towards being increased at divisions 6+ (Figure 3C-D). These results suggest that whereas BCR stimulation merely delays IgG1 CSR to later cell divisions, it directly inhibits IgE CSR.

**Figure 3.**
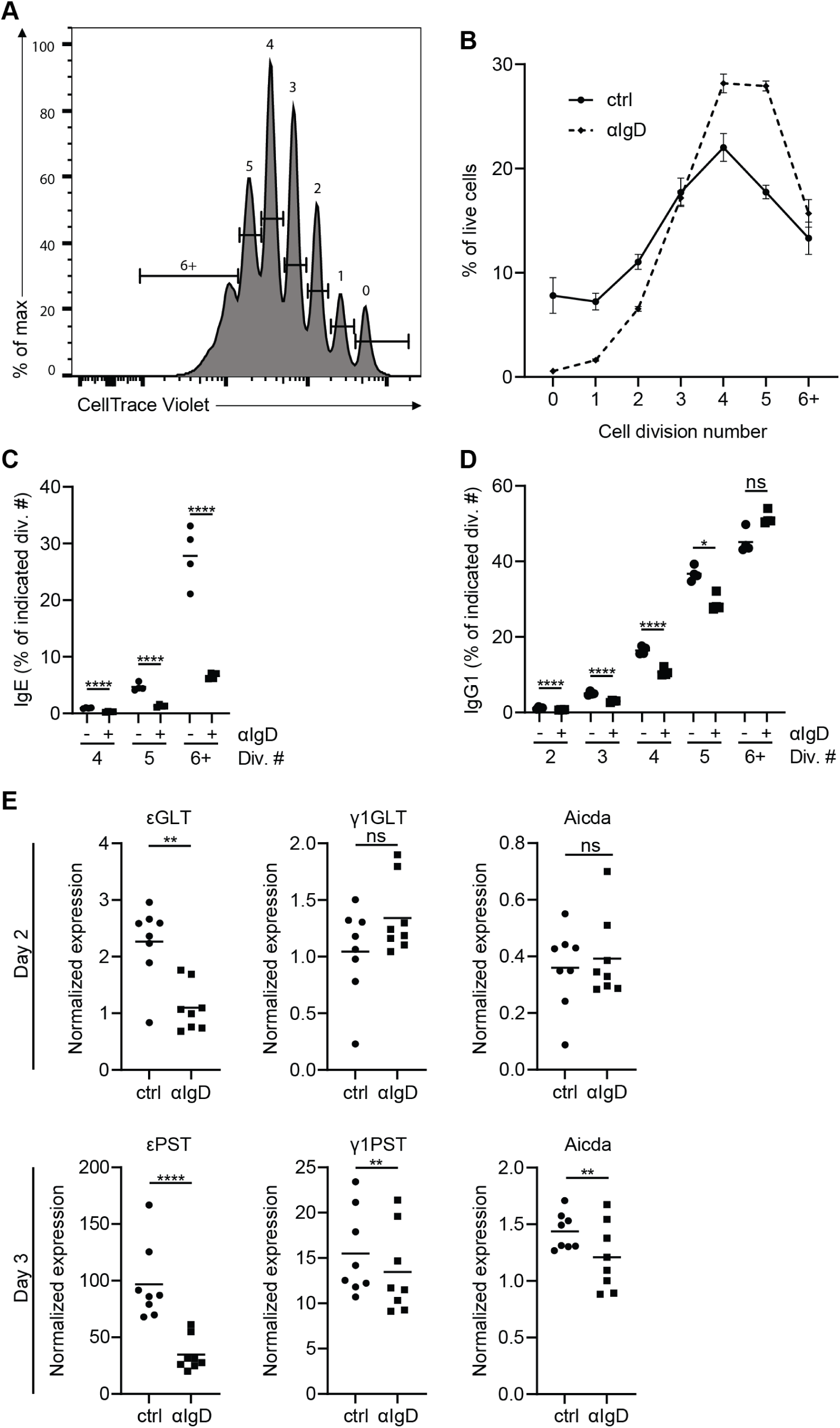
BCR stimulation inhibits IgE CSR. (A-D) Purified naïve mouse B cells were loaded with CTV (see Methods) and then cultured for 4 days with IL-4, αCD40, and control (GGG) or αIgD antibodies (3 μg/mL) prior to analysis by flow cytometry. (A) Representative histogram from the control-treated condition showing CTV staining and cell division number gating. (B) Quantification of the frequency (%) of live cells at each cell division for control- and αIgD-treated conditions, n=4. (C-D) Quantification of the proportion of IgE (C) and IgG1 (D) cells among live cells within each cell division (gated as shown in panel A) for control (black circles) and αIgD (black squares) treatments. (E) Purified naïve mouse B cells were cultured with IL-4, αCD40, and the indicated treatments for two (top row) or three (bottom row) days prior to quantification of the indicated transcripts by RT-qPCR; numbers on the y axis are relative arbitrary units normalized to HPRT. Dots represent average values (B), or samples from individual mice (C-E). Error bars show the SEM (B). Bars represent the mean values (C-E). ns, not significant; *, P < 0.05; **, P < 0.01; ****, P < 0.0001 (one-way repeated measures ANOVA with Dunnett’s post-test comparing the indicated pairs of conditions with the Holm-Sidak correction for multiple comparisons [C-D], paired t test [E]). Results are representative of five similar experiments (A-D) or are pooled from three independent experiments (E).

To gain further insight in the mechanism of IgE CSR inhibition, we quantified ε and γ_1_ germline and post-switch transcripts (GLT and PST, respectively) as well as *Aicda* transcripts (encoding AID) and compared them between BCR-stimulated and control B cell cultures. GLTs are transcribed from the I (e.g. I_ε_ or I_γ1_) region upstream of the switch region for all class-switched antibody isotypes and their production is a pre-requisite for a B cell to undergo CSR to that isotype. It is thought that this is due to their importance in ‘opening up’ local chromatin.^28,29^ The completion of CSR can be detected by the expression of PSTs, which are transcribed from I_μ_ through the recombined switch regions and the downstream constant region. BCR stimulation resulted in a significant >50% reduction in εGLT at D2. In contrast, we observed no significant difference in γ_1_GLT, with a trend toward an increase (28%). In addition, at D3, BCR stimulation led to a strong (64%) reduction in εPST with a subtle (13%) impact on γ_1_PST (Figure 3D). *Aicda* transcripts were equivalent at D2, but at D3 there was a minor (15%) reduction in the αIgD condition. Overall, the more substantial reduction in εPST than γ_1_PST was consistent with the greater impact of BCR ligation on numbers of IgE-switched versus IgG1-switched cells observed above by flow cytometry. Furthermore, the PST data came from D3 of culture, prior to the emergence of most membrane IgE-positive cells, reinforcing our observation in the above CTV experiments that CSR to IgE was reduced at the earliest points at which it could be observed. The reduction in εGLT, but not γ_1_GLT, at D2 is consistent with the notion that the effect of BCR ligation on IgE CSR occurs via altered transcription at the epsilon locus, rather than through changes in *Aicda* transcripts, which we found were unchanged at D2 or reduced only subtly at D3.

### IgE CSR inhibition by BCR stimulation is principally mediated by Syk

As discussed earlier, substantial prior literature has focused on the role of PI3K signaling in CSR.^4,7,16,17,19,20^ Therefore, we set out to resolve whether PI3Kδ, the main PI3K isoform expressed in B cells, is necessary for the inhibition of IgE CSR by BCR stimulation. Consistent with prior work, treatment with the PI3Kδ inhibitor nemiralisib resulted in dose-dependent increases in IgE in the absence of BCR ligation (Figure 4A). However, regardless of the dose of inhibitor, IgE CSR remained strongly susceptible to inhibition by BCR stimulation (Figure 4A). To precisely determine whether PI3Kδ signaling is required for IgE CSR inhibition by BCR stimulation, we performed a normalized analysis. This analysis revealed that the addition of αIgD reduced IgE by ∼70% regardless of the presence of PI3Kδ inhibitor or vehicle control (Figure 4B). These results confirmed that PI3Kδ regulates IgE,^16,20^ but was not required for the inhibition of IgE CSR by BCR stimulation.

**Figure 4.**
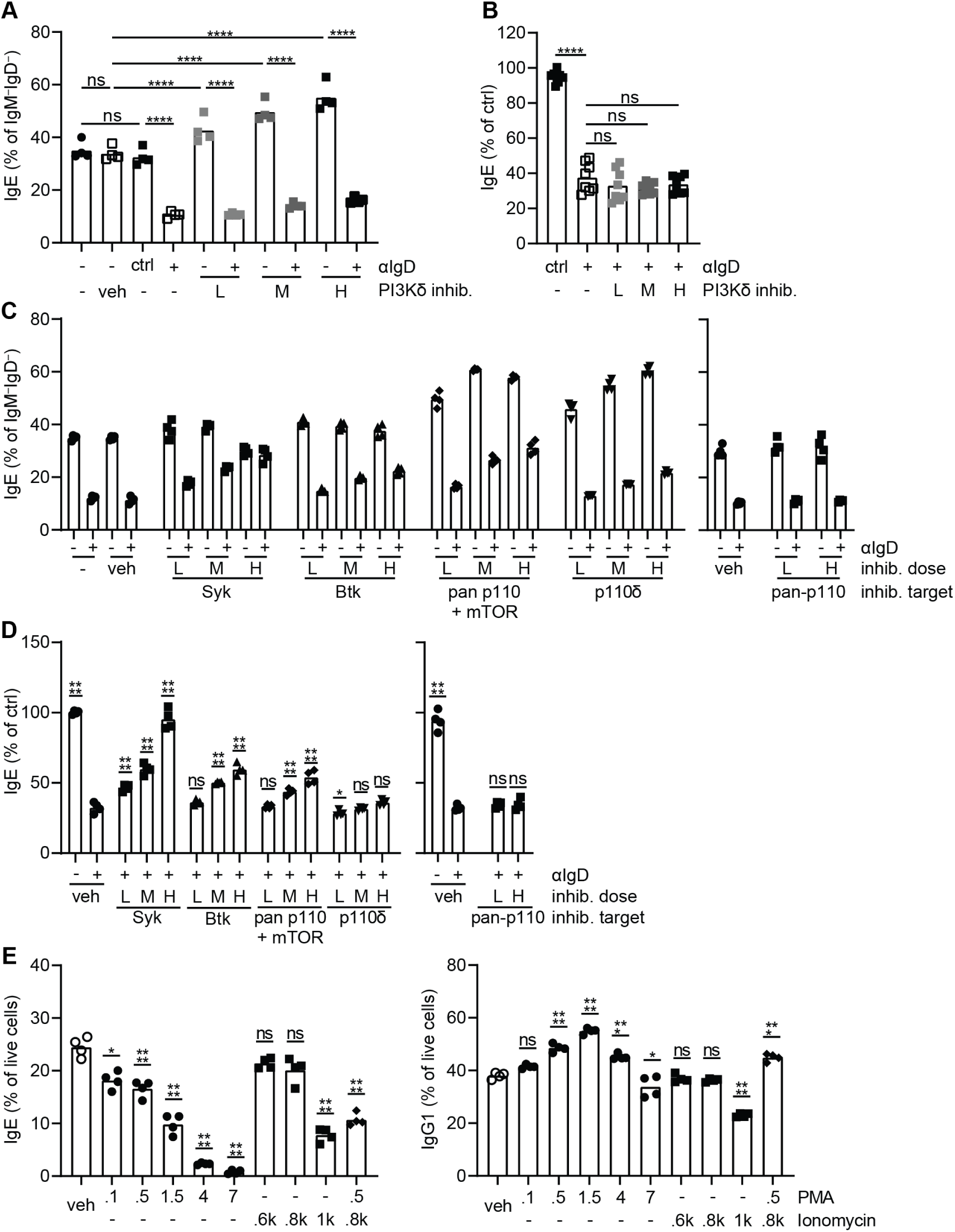
BCR stimulation represses IgE CSR via Syk-dependent rather than p110δ-dependent signaling. (A-E) Purified B cells were cultured with IL-4 and αCD40 for 4 days under various conditions of stimulation and/or inhibitor treatment (as indicated on the x axes) prior to analysis by flow cytometry. (A) Quantification of the proportion of IgE cells among class-switched (IgM^−^IgD^−^) cells according to treatment with αIgD or ctrl (GGG) at 1 µg/mL, as well as different doses of nemiralisib (L – 25nM, M – 100nM, H – 250nM) or vehicle control (DMSO). (B) Normalization was performed by dividing the frequency of IgE cells among class-switched (IgM^−^ IgD^−^) cells in the αIgD-treated (3 μg/mL) condition by their frequency in the control-treated condition (GGG; 1 µg/mL), for each dose of nemiralisib. (C) Quantification of the proportion of IgE cells among class-switched cells treated with αIgD (1 μg/mL; +) or control (no treatment; –), in the presence of vehicle (veh; DMSO) or different doses of inhibitors of Syk (PRT062607, L – 0.4μM, M - 1μM, H – 2.5μM), Btk (ibrutinib; L – 1nM, M – 10nM, H – 50nM), all p110 isoforms and mTOR (omipalisib; L – 1nM, M – 5nM, H – 10nM), p110δ (idelalisib; L – 10nM, M – 50nM, H – 250nM), or all PI3K isoforms (wortmannin; L – 40nM, H – 200nM). See Supplementary Table 1 for statistical comparisons. (D) Quantification, for each dose of inhibitor or vehicle control described for panel D, of the frequency of IgE cells among class-switched (IgM^−^IgD^−^) cells in the αIgD-treated condition as a percentage of their frequency in the untreated control condition. (E) Quantification of the frequency of IgE (left) or IgG1 (right) cells among live cells following treatment with the indicated doses (in ng/mL) of PMA and/or ionomycin. Dots represent samples from individual mice and bars represent the mean values. ns, not significant; *, P < 0.05; **, P < 0.01; ***, P < 0.001; ****, P < 0.0001 (one-way repeated measures ANOVA [A-E] with Dunnett’s post-test comparing the indicated pairs of conditions [A-B], the αIgD-treated condition [D], or the vehicle control [E] using the Holm-Sidak correction for multiple comparisons). See Supplementary Table 1 for statistics for panel C. Results are representative (A, C-E) or pooled from (B) two independent experiments.

We considered various possibilities to explain this result. It could be that an isoform of p110 other than p110δ (e.g. p110α) was responsible for inhibiting IgE CSR downstream of BCR stimulation. Alternatively, there might be redundancy between p110 isoforms, meaning that a complete blockade was required to rescue IgE CSR from inhibition by BCR stimulation. Another possibility is that a non-PI3K-mediated pathway downstream of BCR signaling is responsible for inhibiting IgE CSR. To test these possibilities, we performed further experiments with different inhibitors. To assess if a more complete blockade of p110 signaling could interfere with the inhibition of IgE CSR by BCR stimulation, we selected a second p110δ inhibitor (idelalisib) to validate our earlier findings as well as two pan-p110 inhibitors (omipalisib and wortmannin), one of which also inhibits mTOR (mammalian target of rapamycin; omipalisib). We also tested inhibitors of Syk (PRT062607) and Btk (ibrutinib) to assess an alternative pathway by which BCR stimulation might inhibit IgE CSR. Strikingly, Btk and Syk inhibitors both dose-dependently rescued IgE CSR in the presence of αIgD, but, in the absence of αIgD, left IgE CSR mostly unaffected (Figure 4C). Meanwhile, both idelalisib and omipalisib increased IgE CSR regardless of the presence or absence of αIgD (Figure 4C, see Supplementary Table 1 for statistical comparisons), similar to our earlier results with the PI3Kδ inhibitor. Surprisingly, wortmannin did not increase IgE CSR in the absence of BCR stimulation, unlike the other p110 inhibitors tested. We confirmed the activity of our wortmannin at the relevant dose by verifying its ability to block BCR ligation-dependent S6 phosphorylation (Figure S2). While idelalisib and omipalisib treatment increased IgE CSR, we still observed decreases in IgE CSR following αIgD treatment. To further evaluate if the effects of the p110 inhibitors were independent of BCR stimulation, we again performed a normalized analysis. This analysis confirmed that the highest dose of Syk inhibitor resulted in a complete rescue of IgE CSR from inhibition by BCR stimulation, while maximal Btk inhibition resulted in a substantial yet incomplete rescue (Figure 4D). The p110δ inhibitor idelalisib achieved no significant rescue of IgE CSR inhibition by BCR stimulation, confirming our earlier result. The pan-p110 inhibitor wortmannin also did not rescue IgE CSR, whereas the pan-p110 + mTOR inhibitor omipalisib achieved a moderate rescue. This finding could suggest a contribution of mTOR to IgE CSR inhibition by BCR stimulation. Taken together, these data indicate that PI3K signaling is not required for IgE CSR inhibition by BCR stimulation. Meanwhile, signaling through Syk, with a prominent role for Btk, is required for the inhibition of IgE CSR by BCR stimulation.

Two major outcomes of Syk-dependent BCR signaling are protein kinase C (PKC) activation and Ca^2+^ flux. To examine the sufficiency of these pathways for IgE CSR inhibition we treated cells with phorbol myristate acetate (PMA; a diacylglycerol analog that activates PKC) and/or ionomycin (induces Ca^2+^ flux). PMA treatment led to a strong and dose-dependent reduction in the representation of IgE cells, while, at all but the highest dose tested, IgG1 was either unaffected or increased (Figure 4E). Meanwhile, the highest dose of ionomycin resulted in reductions in both IgE and IgG1, with perhaps a somewhat stronger effect for IgE. The combined effect of PMA and ionomycin together seemed mostly similar to the effect of stimulation with PMA alone. Overall, these data suggest that PKC activation is sufficient to selectively inhibit IgE CSR, whereas Ca^2+^ signaling more broadly inhibits CSR to both IgE and IgG1.

### BCR stimulation acts synergistically with IL-21 or TGFβ1 to inhibit IgE CSR

Having established that BCR signaling can substantially inhibit IgE CSR, we sought to determine if its effects were synergistic with IL-21, which was previously identified as a critical negative regulator of IgE *in vivo*.^2,30^ To this end, we activated B1-8i B cells *in vitro* with IL-4 and αCD40 and treated them with or without cognate antigen and IL-21 (or controls). The combination of cognate antigen and IL-21 resulted in a greater reduction in IgE cells than either alone (Figure 5A, left). Meanwhile, treatment with IL-21 alone or in combination with BCR ligands increased IgG1 cells as a fraction of class-switched cells (Figure 5A, right). These data reveal that BCR stimulation and IL-21 can synergize to inhibit IgE, but not IgG1, CSR.

**Figure 5.**
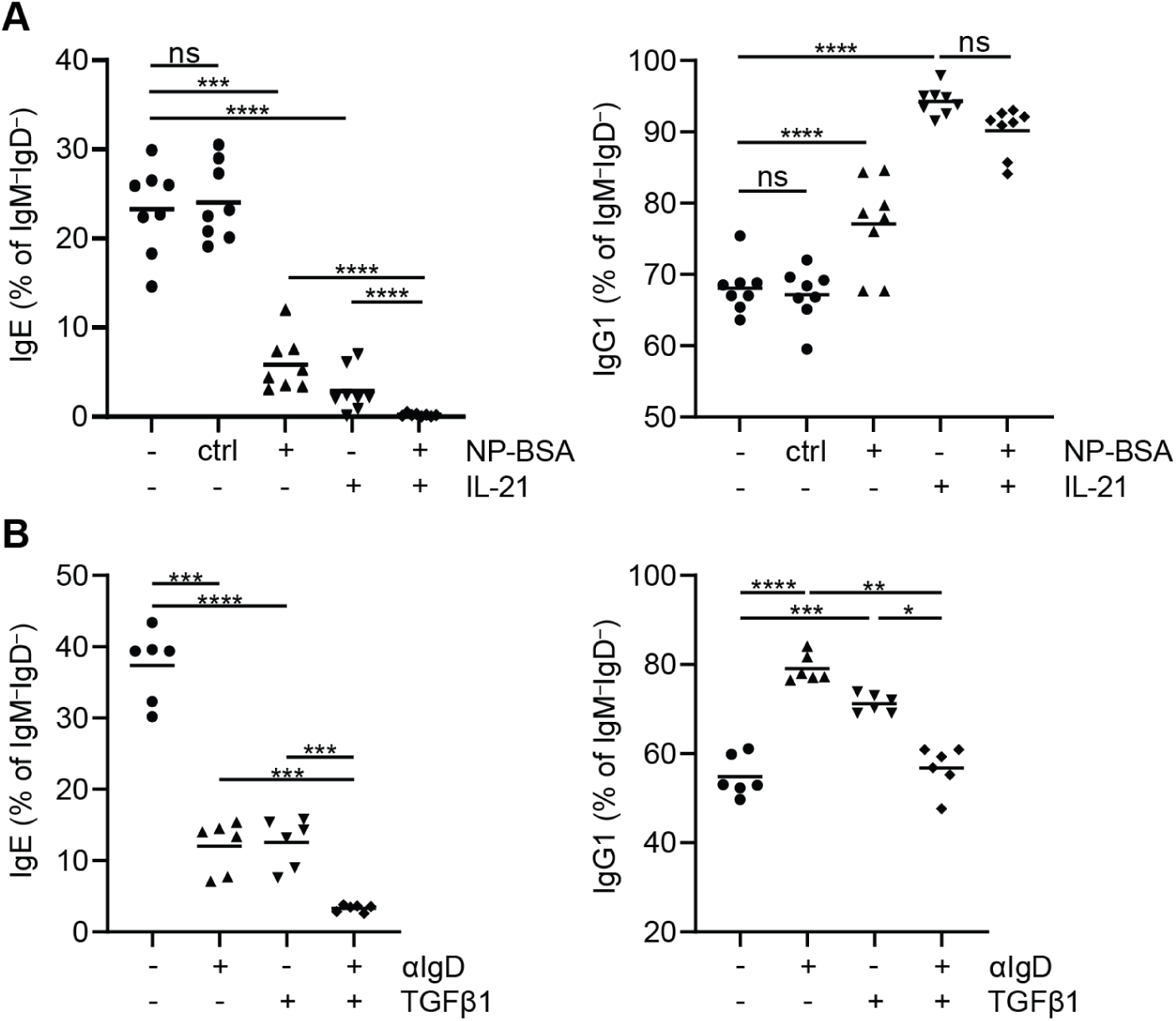
B cell receptor stimulation acts synergistically with IL-21 or TGFβ1 to inhibit IgE CSR. (A) Purified B1-8i B cells were cultured for four days with IL-4, αCD40, and cognate antigen (NP-BSA – 10 ng/mL) or control antigen (ctrl, BSA – 10 ng/mL) in the presence or absence of IL-21 (25 ng/mL). The frequencies of IgE and IgG1 cells among class-switched (IgM^−^IgD^−^) cells were quantified by flow cytometry. (B) Purified B cells were cultured for four days with or without αIgD (3µg/mL) and in the presence or absence of TGFβ1 (2 ng/mL). The frequencies of IgE and IgG1 cells among class-switched cells were quantified by flow cytometry. (A-B) Dots represent samples from individual mice and bars represent the mean. ns, not significant; *, p < 0.05; **, p < 0.01; ***, p < 0.001; ****, P < 0.0001 (one-way repeated measures ANOVA with Dunnett’s post-test comparing the indicated pairs of conditions with the Holm-Sidak correction for multiple comparisons). Results are pooled from two independent experiments.

Having observed synergism between the inhibition of IgE CSR by BCR stimulation and IL-21, we next sought to determine if TGFβ1 possessed similar synergistic capacity. Limited prior evidence suggested that TGFβ1 could inhibit IgE CSR in mouse B cells.^31^ We observed that treatment with TGFβ1 resulted in a dose-dependent reduction in the representation of IgE cells compared to a slight increase in the representation of IgG1 cells (Figure S3A). However, as TGFβ1 is a pleiotropic cytokine with effects on B cell survival and proliferation,^32^ it was unclear whether TGFβ1 could directly inhibit IgE CSR. To resolve this question, as we previously reported for IL-21^2^ and for BCR ligation in Figure 3, we performed experiments with CTV. Culturing B cells with TGFβ1 resulted in strongly reduced proliferation relative to control, evidenced by the substantially larger fraction of cells that did not divide at all and the smaller fraction of cells that divided many times (Figure S3B). Analyzing the rate of IgE and IgG1 switching at each cell division revealed reductions for IgE, but not IgG1, in TGFβ1 treatment conditions, (Figure S3C). Finally, we cultured B cells with αIgD or TGFβ1, alone or in combination, and observed that the combination of both more strongly suppressed IgE than either alone (Figure 5B). These data are consistent with a selective inhibition of IgE CSR by TGFβ1 that is synergistic with BCR stimulation.

### BCR ligation inhibits IgE CSR in human cell culture

Finally, we sought to translate our key findings to humans by investigating the inhibition of IgE CSR by BCR stimulation of tonsillar B cells cultured in conditions that promote CSR to IgE, IgG1, and IgG4. We observed that, similar to our results in mouse cell culture, αIgM treatment resulted in an overall inhibition of CSR (Figure 6A). However, among class-switched cells, αIgM treatment resulted in a dose-dependent reduction in the representation of IgE, but not IgG1 or IgG4, cells (Figure 6B-C). These data are consistent with a selectively enhanced inhibition of IgE CSR relative to IgG1 or IgG4. Next, we investigated the synergism between BCR stimulation and IL-21 treatment of human B cells and found that, as with mouse B cells, BCR ligation and IL-21 together led to a greater reduction in IgE than either alone (Figure 6D). αIgM treatment and/or IL-21 treatment did not clearly affect IgG4, whereas the representation of IgG1 cells was enhanced by IL-21 in a manner which was not affected by the presence/absence of αIgM. These findings support the notion that BCR stimulation is a conserved, selective inhibitor of IgE CSR in mice and humans.

**Figure 6.**
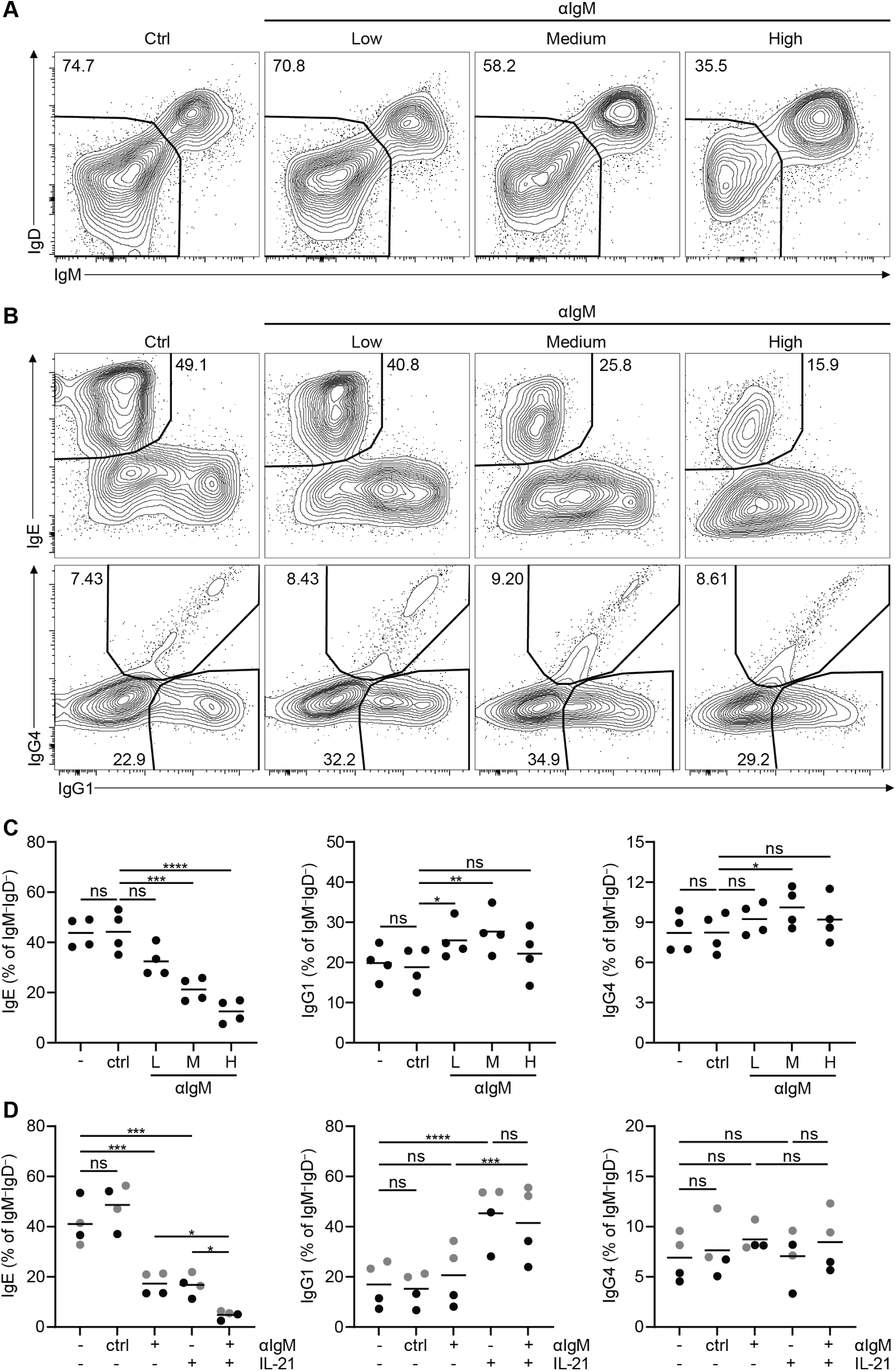
BCR stimulation inhibits IgE CSR in human B cells. (A-D) Purified human tonsillar naïve B cells were cultured in IL-4, IL-10, αCD40, and the indicated treatment conditions for 8 days prior to analysis by flow cytometry. (A-B) Representative flow cytometry plots and gating strategies for class-switched (IgM^−^IgD^−^) cells (A), IgE cells (B; top row), or IgG1 and IgG4 cells (B; bottom row) for cells treated with low, medium, or high (0.1, 0.3, or 3 µg/mL, respectively) doses of αIgM or control (GGG; 3 µg/mL). Cells in (B) were pre-gated as IgM^−^IgD^−^, as shown in (A). Note that the IgG1 antibody cross-reacts with IgG4 and therefore IgG4-expressing cells appear as IgG4^+^ IgG1^+^, whereas IgG1-expressing cells appear as IgG4^−^ IgG1^+^. (C-D) The proportions of IgE (left), IgG1 (center), and IgG4 (right) cells within the class-switched (IgM^−^IgD^−^) compartment were quantified by flow cytometry according to the indicated treatment conditions. (C) Quantification of the impact of αIgM titration on CSR. Treatment doses were as described for panels A-B. (D) Quantification of the synergistic impacts of IL-21 and BCR ligation on CSR. Treatment doses were: ‘-’ – untreated; ctrl – GGG at 300 ng/mL (grey) or 2 µg/mL (black), αIgM – 300 ng/mL (grey) or 2 µg/mL (black); IL-21 – 5 (grey) or 10 (black) ng/mL. (C-D) Dots represent samples from individual human tonsil donors and bars represent the mean. ns, not significant; *, P < 0.05; **, P < 0.01; ***, P < 0.001; ****, P < 0.0001 (one-way repeated measures ANOVA with Dunnett’s post-test comparing the indicated pairs of conditions and the Holm-Sidak correction for multiple comparisons). Results are representative of (A-B) or pooled from (C-D) two experiments.

## Discussion

Here, we report that BCR stimulation selectively inhibits CSR to IgE in mouse and human B cells. In mice, the representation of IgE cells in PC and GC compartments was inversely regulated by BCR affinity and surface expression. We relate these findings to the inhibition of IgE CSR by finding reduced εGLT and fewer IgE-switched cells across cell divisions in cells stimulated through their BCRs versus those left untreated. A major finding is that the selective inhibition of IgE CSR by BCR signaling required Syk, rather than PI3K as had been proposed, and could be mimicked by the activation of PKC. BCR stimulation synergized with IL-21 or TGFβ1 to inhibit IgE CSR more greatly than with any one stimulus alone. Finally, we replicated our findings from murine studies of selective inhibition of IgE CSR by BCR stimulation as well as synergy for IgE CSR inhibition by IL-21 and BCR stimulation in cultures of human tonsillar B cells. These observations establish that IgE CSR is uniquely susceptible to inhibition by BCR signaling.

The present work extends our understanding of the inhibition of IgE CSR by BCR stimulation. Prior work left unclear whether BCR stimulation had a broad effect on CSR^6,7^ or a selective effect on IgE^11^. Furthermore, while prior studies exclusively used antibodies to stimulate the BCR,^5–7,33^ we additionally included data with cognate antigen *in vitro* and *in vivo*. These experiments allowed us to examine if different cognate antigens exhibited different selectivity for IgE versus IgG1 CSR inhibition. While all variants we tested resulted in greater inhibition of IgE than IgG1 CSR, some intermediate doses exhibited exquisite specificity for IgE whereas high doses could strongly reduce both IgE and IgG1. Interestingly, low doses of high-affinity high-valency antigen actually resulted in increased IgG1 CSR relative to control, which may relate to a prior report that identified increased IgG1 switching in LPS culture with high-affinity antigen.^34^ A similar finding was also reported for IgA.^35^ Meanwhile, we identified no conditions under which BCR ligation resulted in increased IgE, reinforcing our observations of selective IgE CSR inhibition by BCR stimulation.

Our qPCR results indicate that a likely mechanism for the selective inhibition of IgE CSR by BCR stimulation is a reduction in εGLT. This finding contrasts with earlier work exploring the mechanism by which BCR stimulation inhibited CSR, which did not observe effects on εGLT.^6^ One key difference in our studies was in the timepoint analyzed: we measured εGLT at day 2, which we have found is a key timepoint prior to CSR,^2^ whereas Jabara *et al.*^6^ analyzed εGLT at day 4, a timepoint at which we have found CSR has already occurred. Our quantitative analysis may also have been more sensitive in detecting the reduction in εGLT. Rather than εGLT, prior work focused on a temporary, BCR stimulation-induced reduction in *Aicda* expression as a potential mechanism for CSR inhibition.^6,7^ However, a series of studies from a different group^5,33^ found that, while BCR stimulation reduced AID, AID overexpression actually reduced IgE CSR. Overall, there is no consistent link between changes in AID protein or gene expression and IgE CSR following BCR ligation. Here, we found that *Aicda* transcripts were unchanged relative to control at day 2 after BCR stimulation and were slightly reduced at day 3. For IgE, the more substantial and earlier BCR ligation-induced reduction in εGLT relative to *Aicda*, in addition to aforementioned evidence^5,33^, indicates that differences in εGLT levels represent a more plausible explanation for strongly reduced CSR. Relating to the broad effects of BCR signaling on CSR under some conditions, our findings do not exclude a role for alterations in AID gene or protein expression.

Using various inhibitors, we establish a Syk-dependent rather than p110δ-dependent signaling mechanism for the inhibition of IgE CSR by BCR stimulation. Our observations of the lack of a strong rescue of IgE CSR with the pan-p110/mTOR inhibitor omipalisib, or the lack of any rescue with the pan-p110 inhibitor wortmannin, in the face of BCR stimulation were unexpected given a prior report that a PI3Kα/δ/β inhibitor completely rescued IgG1 CSR from inhibition by BCR stimulation.^7^ This difference in the extent of CSR rescue for IgE and IgG1 might indicate that BCR-dependent effects on IgG1 CSR are moreso driven by PI3K signaling, whereas we have shown that BCR-dependent effects on IgE CSR require Syk-dependent signaling pathways. This model might also help to explain the selectively enhanced impact of BCR stimulation on IgE switching relative to IgG1. Alternatively, it could be that a maximal blockade of PI3K is required to observe an effect on BCR ligation-induced IgE CSR inhibition, and that this maximal blockade cannot be achieved with chemical inhibitors due to the necessity of PI3K signaling for B cell survival.^36^ However, at least for wortmannin, this possibility seems less likely as we found that 200nM (a dose comparable to that previously observed to effectively inhibit primary mouse B cell chemotaxis)^37^ was sufficient to completely block detectable BCR ligation-induced S6 phosphorylation, consistent with a complete blockade of PI3K activity (Figure S2). In addition to our studies with inhibitors, we used PMA and ionomycin to bypass the BCR and demonstrate that PKC activation, and, to a lesser extent, Ca^2+^ signaling, were sufficient to inhibit IgE CSR, with PKC activation having a more selective effect on IgE. Overall, we find greater evidence that the inhibition of IgE CSR depends on a Syk-Btk signaling axis than PI3K.

While we did not find a role for PI3K signaling in the inhibition of IgE CSR by antigen-induced BCR signaling, several PI3K inhibitors had clear impacts on baseline IgE switching, consistent with prior reports.^16,20^ The mechanism underlying the impact of these inhibitors on IgE CSR remains unclear, including whether the inhibitors are modifying PI3K activity related to antigen-independent (tonic) BCR signaling, such as that which is required for B cell survival, or PI3K signaling downstream of other pathways entirely. Interestingly, observations of p110 inhibitors increasing baseline switching to IgE in murine cell culture may not translate to humans, as inhibition of p110δ was reported to reduce, rather than increase, IgE in cultured human B cells.^38^ If confirmed, this would suggest that PI3K activity might differently regulate IgE CSR in non-BCR stimulated human versus mouse B cells.

Our finding of synergy between BCR signaling and IL-21 for the inhibition of IgE CSR may be particularly important *in vivo*. In a physiologic response, B cells possess a variety of starting affinities for cognate antigen, and multiple antigens may be present in varying amounts. Our data suggests that lower-affinity B cells, or B cells responding to a scarcer antigen, may be relatively advantaged for IgE CSR, as they would experience weaker or fewer BCR signaling events relative to high-affinity cells responding to abundant antigen. Varying antigen encounters would also have ramifications for subsequent T:B interactions, as B cells that bind greater amounts of antigen would not only have greater BCR signaling but also greater antigen capture. Greater antigen capture, leading to more antigen presentation, could potentially influencing the nature of help received from T follicular helper cells (T_FH_), which are notably important sources of IL-21 and IL-4.^1,39^ Therefore, our findings imply that IgE CSR may be more likely to occur in lower-affinity B cells engaged with T_FH_ producing less IL-21 but ample IL-4.^1,40^ We also found that TGFβ1 treatment inhibited IgE CSR and could synergize with BCR ligation. Importantly, we found that, although TGFβ1 globally limited B cell proliferation, it also resulted in a selective reduction in IgE at each cell division, consistent with IgE CSR inhibition. The timing of TGFβ1 signaling related to B cell activation *in vivo* is perhaps less clear than for IL-21 or BCR ligation, but activated B cells have been described to undergo autocrine TGFβ1 signaling critical for maintaining immune tolerance,^41^ indicating that B cell activation is associated with TGFβ1 signaling that could restrict IgE CSR.

Our finding that BCR stimulation also impaired human IgE CSR provides an important piece of translational evidence for this topic, which until now was restricted to mouse studies. We found that αIgM resulted in similar inhibition of IgE CSR in human cells to that which we observed in mouse cells. It will be interesting to determine in the future if the inhibition of IgE CSR by BCR stimulation in human cells is Syk-dependent, as we observed in mouse B cells. Notably, there are clear links between human mutations that impair antigen-receptor signaling and elevated IgE or atopy.^42^ However, in addition to potential impacts on B cell CSR, alterations in antigen receptor signal transduction could affect T cells, IgE B cells,^8,10^ and/or IgE PCs;^3^ therefore, it will be important to disentangle the contributions made by each cell type to allergic disease pathogenesis.

This study defines ligand-induced BCR signaling as a selective, negative regulator of IgE CSR in mouse and human B cells. Our finding that BCR signaling and specific cytokines (IL-21 or TGFβ1) synergized to inhibit IgE CSR reveals how multiple layers of regulation can cooperate to suppress the development of allergic immunity. This raises the question of whether the development of allergy is associated with signaling impairments downstream of the BCR and/or cytokine receptors that normally inhibit IgE CSR. Overall, we believe that our findings are of significance for understanding the homeostatic regulation of allergic immunity, with implications for allergic disease pathogenesis.

## Methods

### Mice and immunizations

All mice used for experiments in this study were on a C57BL/6 (B6) background (backcrossed ≥10 generations). Mice for experiments were sex and age-matched between groups as much as possible and both male and female mice were used. For *in vivo* experimentation, mice were at least 6 weeks of age and for *in vitro* experimentation donor animals were at least 5 weeks of age. Mice were housed in specific-pathogen-free facilities. Mouse work was approved by the Institutional Animal Care and Use Committee (IACUC) of the University of California, San Francisco (UCSF).

B1-8i (012642; B6.129P2(C)-*Igh*^tm2Cgn^/J), Boy/J (002014; B6.SJL-*Ptprc*^a^*Pepc*^b^/BoyJ), B6/J (000664; C57BL/6J), and B6 Thy1.1 (000406; B6.PL-*Thy1^a^*/CyJ) mice were originally from The Jackson Laboratory and were maintained in our colony. B6/J, Boy/J, and B6 Thy1.1 mice were used as “WT” throughout. Immunizations consisted of antigen dissolved in D-PBS and mixed 50:50 volumetrically with alum (Alhydrogel; Accurate Chemical and Scientific) injected in a volume of 20µL subcutaneously into the ear pinnae. For immunization of Hy10 recipients in Figure 1A, 6.25µg DEL-OVA or HEL-OVA was used (see *HEL-OVA/DEL-OVA preparation* below). For immunization of wildtype and Igα^+/-^ mice in Figure 1B-C, 10µg NP conjugated to chicken gamma globulin (NP-CGG; Biosearch Technologies) was used. The left and right facial LNs were pooled for analysis at endpoint (d7) by flow cytometry.

### HEL-OVA/DEL-OVA preparation

HEL/DEL-OVA conjugations were carried out as using maleimide thiol chemistry as described previously.^10^ Correct product formation was verified with SDS-PAGE. Finally, products were separated from reactants by FPLC, enrichment was assessed using SDS-PAGE, and concentration was determined by A_280_.

### Mouse cell culture, in vitro BCR stimulation, and inhibitor treatments

All *in vitro* mouse cell culture experiments (except phosflow experiments, see below) were performed with either total (Figures 1, 3, 4, and 5B), or Igκ-negative (Figures 2 and 5A) B cells, purified by negative selection as described previously.^3^ After purification, cells were resuspended in complete RPMI (cRPMI), composed of RPMI 1640 without L-glutamine (Thermo Fisher Scientific), 10% FBS, 10 mM Hepes, 1X penicillin streptomycin L-glutamine (Thermo Fisher Scientific), and 50 μM β-mercaptoethanol (Thermo Fisher Scientific. Purified B cells were seeded at a density of 2-10 * 10^3^ cells per well in 96-well Microtest U-bottom plates (BD Falcon) and were cultured with αCD40 (150 ng/mL; clone FGK-45; Miltenyi Biotec) and IL-4 (25 ng/mL; Peprotech) for prior to analysis by flow cytometry (d4) or RT-qPCR (d2-3). Cells were plated in triplicate for each condition, except for some CTV experiments where sextuplicates were used. In some cases, CD45-congenic B cells were co-plated to allow the combined assessment of cells from two different mice.

BCR stimulation with antibodies was performed using goat anti-mouse IgD (Nordic MUbio), goat polyclonal F(ab’)2 anti-mouse Igκ (LifeSpan Biosciences), goat polyclonal F(ab’)2 anti-mouse IgM (αIgM; Jackson ImmunoResearch), or Chrompure Goat IgG (Jackson ImmunoResearch) as a control. BCR stimulation with cognate antigen was performed using NP(4)BSA (Biosearch Technologies), NP(25)BSA (Biosearch Technologies), NIP_24_BSA (Biosearch Technologies), or BSA (Sigma-Aldrich) as a control. Recombinant human TGFβ1 (Peprotech) was used for experiments in Figure 5. Stimulations were prepared in the culture medium at the doses indicated in figure legends. All inhibitors were diluted in DMSO. The concentration of DMSO in culture never exceeded 0.1% and was typically lower. The specific DMSO vehicle control concentration in each experiment was made to be equivalent to the highest concentration of DMSO in any inhibitor-treated well. See figure captions for specific inhibitors and concentrations used.

### CellTrace Violet labeling

Labeling with CellTrace Violet (CTV; Life Technologies Corporation) was performed as described previously.^2^ Briefly, cells were incubated with CTV at 1μM for 20 minutes in a 37°C water bath and then washed twice with FBS (Life Technologies Corporation) underlaid in each wash step. Pilot experiments revealed that CTV labeling and associated centrifugation steps resulted in a ∼75% reduction in cell number, which was taken into account when plating cells after labeling.

### RNA extraction, cDNA conversion, and RT-qPCR amplification

Harvesting of nucleic acids, the preparation of cDNA, and RT-qPCR amplification, standard curve preparation, and analysis were performed as described previously.^2^ The following specific primers were used for the qPCR assays:

ε GLT forward: 5’-TCGAATAAGAACAGTCTGGCC-3’ ε GLT reverse: 5’-TCACAGGACCAGGGAAGTAG-3’

γ1 GLT forward: 5’-CAGGTTGAGAGAACCAAGGAAG-3’

γ1 GLT reverse: 5’-AGGGTCACCATGGAGTTAGT-3’

*Aicda* forward: 5’ CCTAAGACTTTGAGGGAGTCAA-3’

*Aicda* reverse: 5’-CACGTAGCAGAGGTAGGTCTC-3’

*Hprt* (internal control) forward: 5’-TGACACTGGCAAAACAATGCA-3’

*Hprt* (internal control) reverse: 5’GGTCCTTTTCACCAGCAAGCT-3’

### Human samples, B cell purification, and cell culture

Human B cells were purified from tonsils fractions obtained from UCSF pathology. Excess tissue collected following routine tonsillectomies was completely de-identified, allowing samples to not be classified as human subjects research according to the guidelines from the UCSF Institutional Review Board. Tonsillar tissue was dissociated, a single cell suspension was prepared and then cells were cryopreserved as described previously.^43^ Naive B cells were purified by magnetic bead depletion using the Mojosort Human Naive B Cell Isolation kit (BioLegend) according to manufacturer’s instructions but with some additional modifications as previously described.^2^ Following purifications, naïve human B cells were resuspended in complete Iscove’s modified Dulbecco’s medium (cIMDM), consisting of: IMDM supplemented with GlutaMAX^TM^ (Gibco), 10 % FBS, 1X pencillin-streptomycin (UCSF Cell Culture Facility), 1X insulin-transferrin-selenium (ITS-G, Fiser Scientific), 0.25 μg/mL Amphotericin B (Neta Scientific), and 100 IU/mL Nystatin (Neta Scientific).

A fraction of purified cells was analyzed by flow cytometry to verify purity. Cells were then cultured at a density of 5-20k live B cells/well in 96-well Microtest U-bottom plates (BD Falcon)for 8 days with 100 ng/mL anti-human CD40 antibody (clone G28.5; Bio-X-Cell), 25 ng/mL recombinant human IL-4 (Peprotech), and 50 ng/mL human IL-10 (Peprotech). Where indicated in the figure caption, 50 ng/mL human IL-21 (Peprotech), goat anti-human IgM F(ab’)2 fragments, or GGG was added in addition.

### Flow cytometry

Processing of dLNs and downstream flow cytometric analysis was performed as previously described.^3^ For both *in vivo* and *in vitro* experiments all incubations were 20 minutes on ice except for Fc block incubations (10 minutes) and antibody staining of fixed and permeabilized cells (45 minutes to 1 hour). For human experiments, excess mouse gamma globulin was used to block, as anti-human fluorochrome conjugates were mouse IgG antibodies. See Supplementary Tables 2 and 3 for mouse and human flow cytometry reagents, respectively.

For both mouse and human experiments, we used our previously-established intracellular staining technique^9^ to sensitively and specifically detect IgE-expressing cells. Briefly, to prevent the detection of IgE captured by non-IgE-expressing cells, surface IgE was blocked with a large excess of unconjugated αIgE (clone RME-1 for mouse experiments, clone MHE-18 for human experiments). IgE-expressing cells were then detected after fixation/permeabilization by staining with a low concentration of fluorescently-labelled αIgE of the same clone.

After staining, cells were collected on an LSRFortessa (BD). Data were analyzed using FlowJo v10. Counting beads were identified by their high SSC and extreme fluorescence and were used to determine the proportion of the cells plated for staining that had been collected on the flow cytometer for each sample. Cells were gated on FSC-W versus FSC-H and then SSC-W versus SSC-H gates to exclude doublets, and next as negative for the fixable viability dye eFluor780 and over a broad range of FSC-A to capture resting and blasting live lymphocytes. 2D plots were presented as contour plots with outliers shown as dots.

### Phosflow

One million splenocytes in 200µL cRPMI/well were plated in a 96-well Microtest U-bottom plates (BD Falcon). Different conditions were plated in duplicate. Wortmannin or an equivalent volume of vehicle control (DMSO) was added to a final concentration of 200nM. Cells were then incubated at 37°C for 10 minutes in a 5% CO_2_ incubator to allow inhibitor binding. Cells were pelleted at 730 × *g* and then resuspended using 25µL of either control or αIgD-containing solution, each of which also contained fixable viability dye eFluor780 at a 1/500 dilution. Cells were returned to the incubator for a further 10 minutes to allow for BCR signaling-dependent phosphorylation events and viability staining. After this incubation, 100µL of Phosflow Fix Buffer 1 (BD Biosciences), pre-warmed to 37°C, was added to each well and cells were incubated for a further 11 minutes. Cells were pelleted at 930 × *g*, washed twice with FACS buffer, then stained for surface markers on ice as normal. Following surface staining, cells were washed then permeabilized with 200µL/well Phosflow Perm/Wash Buffer 1 (BD Biosciences) for 23 minutes at room temperature. Cells were then pelleted at 930 × *g* and resuspended in 25µL of Phosflow Perm/Wash Buffer 1 containing 1/5-diluted αpS6-AF647 and incubated at room temperature for 1 hour. After staining, cells were washed and resuspended in FACS buffer prior to analysis by flow cytometry.

### Statistical analysis

To achieve power to discern meaningful differences, experiments were performed with multiple biological replicates and/or multiple times, see figure legends. The number of samples chosen for each comparison was determined based on past similar experiments to gauge the expected magnitude of differences. GraphPad Prism v9 was used for statistical analyses. Data approximated a log-normal distribution and thus were log transformed for statistical tests. Statistical tests were selected by consulting the GraphPad Statistics Guide according to experimental design. All tests were two-tailed. Groups were assumed to have similar standard deviation for ANOVA analysis.

## Figures

**Figure S1.**
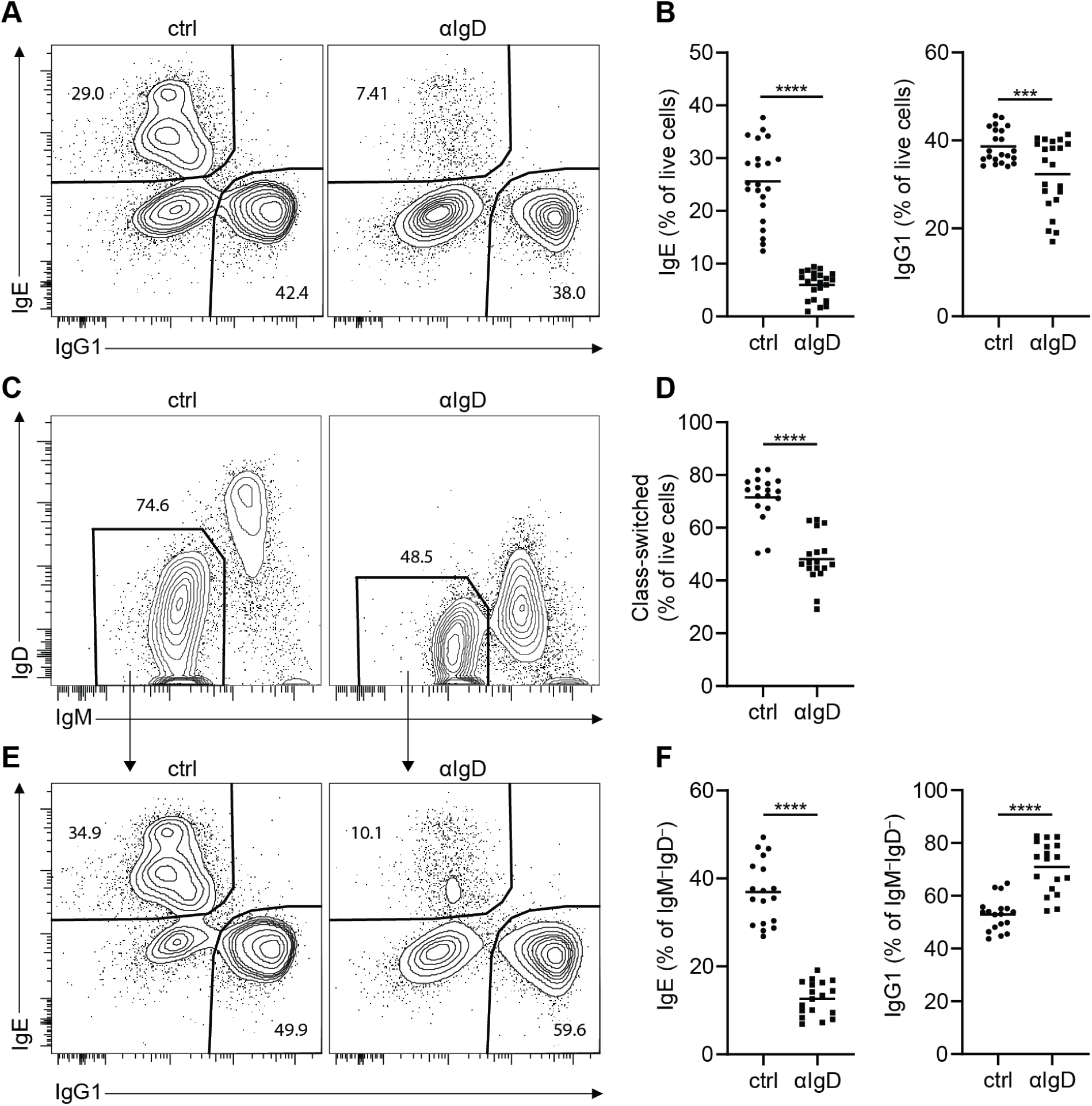
Supporting data for Figure 1. (A-F) B cells were cultured with IL-4 and αCD40 for 4 days prior to analysis by flow cytometry with control (GGG, 3 μg/mL) or αIgD (3 µg/mL) antibodies. (A-B) Representative flow cytometry plots (A) and quantification (B) of the impact of BCR ligation on the frequency of IgE and IgG1 cells among live cells. (C-D) Representative flow cytometry plots (C) and quantification (D) of the impact of BCR ligation on the frequency of class-switched (IgM^−^IgD^−^) cells among live cells. (E-F) Representative flow cytometry plots (E) and quantification (F) of the impact of BCR ligation on the frequency of IgE and IgG1 cells among class-switched (IgM^−^IgD^−^) cells. (B, D, F) Dots represent samples from individual mice and bars represent the mean values. ***, P < 0.001; ****, P < 0.0001 (paired t test). Results are representative of six (A) or five (C, E), or are pooled from six (B) or five (D, F), independent experiments.

**Figure S2.**
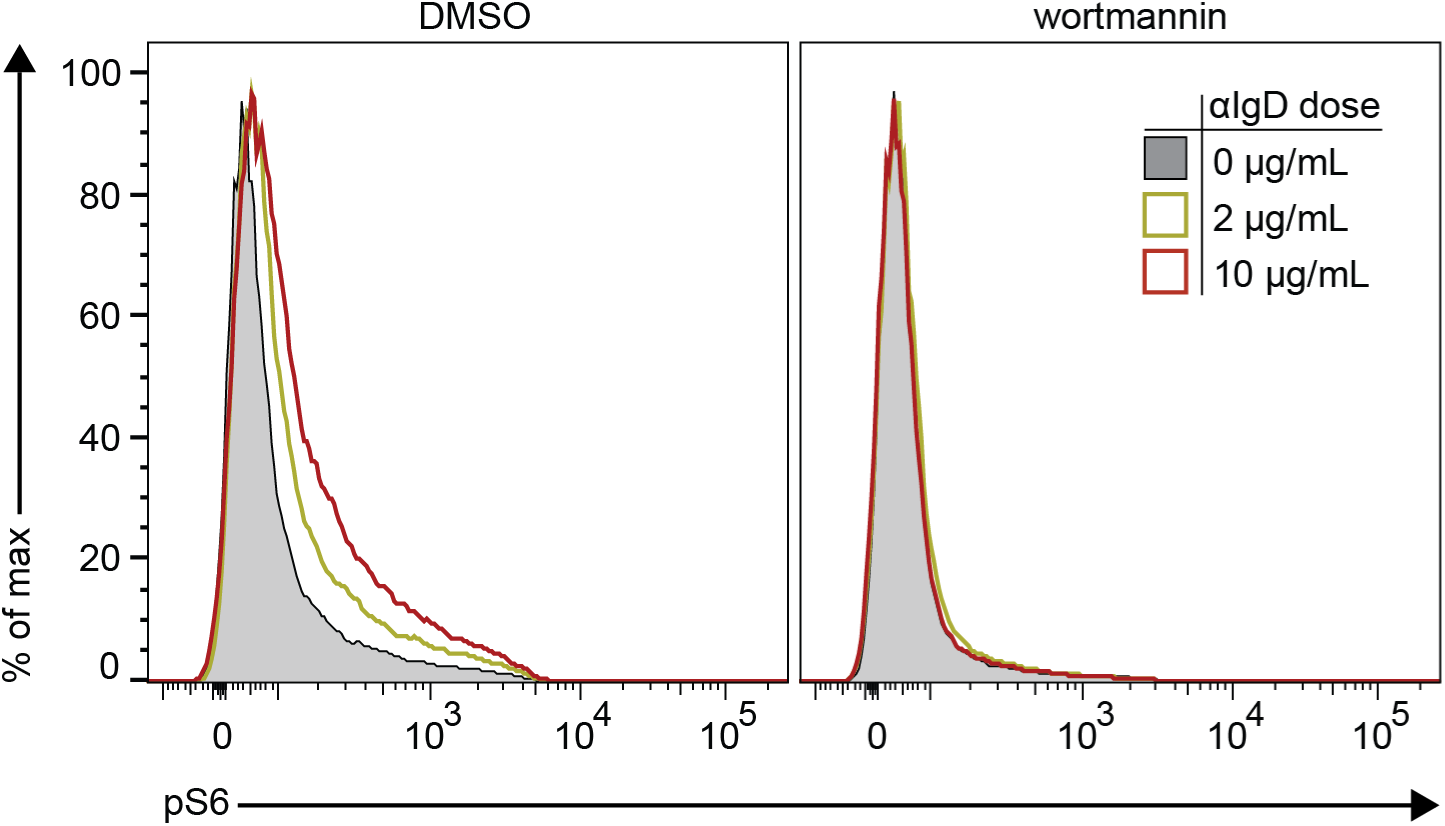
Supporting data for Figure 4. Cultured B cells were treated with the indicated dose of αIgD and with either 200nM wortmannin or DMSO vehicle control prior to analysis of phosphorylated S6 (pS6) by phosflow (see Methods). Data are representative of two independent experiments.

**Figure S3.**
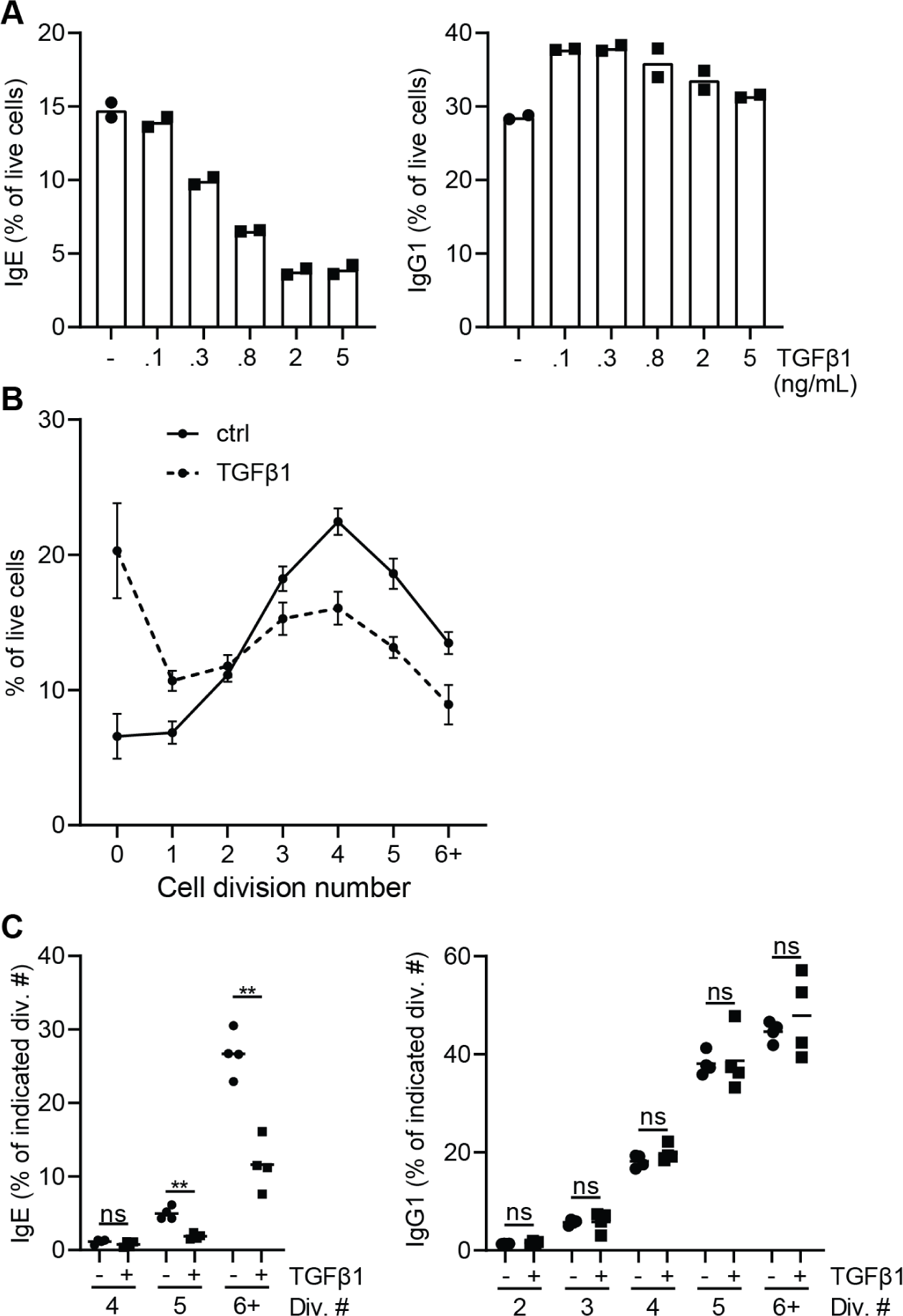
TGFβ1 inhibits IgE class switching. (A-C) Purified naïve B cells were loaded with CTV (see Methods) and then cultured for 4 days with IL-4, αCD40, and vehicle control (DMSO) or TGFβ1 prior to analysis by flow cytometry. (A) The rate of switching to IgE (left) or IgG1 (right) in B cell culture according to TGFβ1 dose. (B) Quantification of the percentage of live cells at each cell division for control- and TGFβ1-treated conditions, n=4. (C) Quantification, for each of the indicated cell divisions, of the frequency of IgE (left) and IgG1 (right) cells among live cells, according to treatment group (control, black circles; αIgD, black squares). TGFβ1 doses were 0 or 2 ng/mL (– or +, respectively). (A,C) Dots represent samples from individual mice and bars represent the mean. (B) Dots represent the mean, errors bars show the SEM. ns, not significant; **, p<0.01 (one-way repeated measures ANOVA with Dunnett’s post-test comparing the indicated pairs of conditions with the Holm-Sidak correction for multiple comparisons). Except for panel A, results are representative of two independent experiments.

**Supplementary Table 1.** Šídák’s multiple comparisons testing of repeated measures ANOVA.

| **Left side of Figure 4C** | **Summary** | **Adjusted P Value** |
| --- | --- | --- |
| neg vs. neg + αIgD | **** | <0.0001 |
| veh vs. veh + αIgD | **** | <0.0001 |
| Ibrutinib (50nM) vs. ibrutinib (50nM) + αIgD | **** | <0.0001 |
| Ibrutinib (10nM) vs. ibrutinib (10nM) + αIgD | **** | <0.0001 |
| Ibrutinib (1nM) vs. ibrutinib (1nM) + αIgD | **** | <0.0001 |
| PRT062607 (4uM) vs. PRT062607 (4uM) + αIgD | ns | >0.9999 |
| PRT062607 (2.5uM) vs. PRT062607 (2.5uM) + αIgD | ns | >0.9999 |
| PRT062607 (1uM) vs. PRT062607 (1uM) + αIgD | **** | <0.0001 |
| PRT062607 (.4uM) vs. PRT062607 (.4uM) + αIgD | **** | <0.0001 |
| Omipalisib (10nM) vs. omipalisib (10nM) + αIgD | **** | <0.0001 |
| Omipalisib (5nM) vs. omipalisib (5nM) + αIgD | **** | <0.0001 |
| Omipalisib (1nM) vs. omipalisib (1nM) + αIgD | **** | <0.0001 |
| Idelalisib (250nM) vs. idelalisib (250nM) + αIgD | **** | <0.0001 |
| Idelalisib (5nM) vs. idelalisib (5nM) + αIgD | **** | <0.0001 |
| Idelalisib (10nM) vs. idelalisib (10nM) + αIgD | **** | <0.0001 |
| neg vs. veh | ns | >0.9999 |
| veh vs. ibrutinib (50nM) | ns | 0.9968 |
| veh vs. ibrutinib (10nM) | ns | 0.416 |
| veh vs. ibrutinib (1nM) | * | 0.0455 |
| veh vs. PRT062607 (4uM) | **** | <0.0001 |
| veh vs. PRT062607 (2.5uM) | ns | 0.0671 |
| veh vs. PRT062607 (1uM) | ns | 0.5302 |
| veh vs. PRT062607 (.4uM) | ns | 0.9692 |
| veh vs. omipalisib (10nM) | **** | <0.0001 |
| veh vs. omipalisib (5nM) | **** | <0.0001 |
| veh vs. omipalisib (1nM) | **** | <0.0001 |
| veh vs. idelalisib (250nM) | **** | <0.0001 |
| veh vs. idelalisib (5nM) | **** | <0.0001 |
| veh vs. idelalisib (10nM) | **** | <0.0001 |
| neg + αIgD vs. veh + αIgD | ns | 0.9966 |
| neg + αIgD vs. ibrutinib (50nM) + αIgD | **** | <0.0001 |
| neg + αIgD vs. ibrutinib (10nM) + αIgD | **** | <0.0001 |
| neg + αIgD vs. ibrutinib (1nM) + αIgD | ** | 0.0023 |
| neg + αIgD vs. PRT062607 (4uM) + αIgD | **** | <0.0001 |
| neg + αIgD vs. PRT062607 (2.5uM) + αIgD | **** | <0.0001 |
| neg + αIgD vs. PRT062607 (1uM) + αIgD | **** | <0.0001 |
| neg + αIgD vs. PRT062607 (.4uM) + αIgD | **** | <0.0001 |
| neg + αIgD vs. omipalisib (10nM) + αIgD | **** | <0.0001 |
| neg + αIgD vs. omipalisib (5nM) + αIgD | **** | <0.0001 |
| neg + αIgD vs. omipalisib (1nM) + αIgD | **** | <0.0001 |
| neg + αIgD vs. idelalisib (250nM) + αIgD | **** | <0.0001 |
| neg + αIgD vs. idelalisib (5nM) + αIgD | **** | <0.0001 |
| neg + αIgD vs. idelalisib (10nM) + αIgD | ns | 0.9994 |

| **Right side of Figure 4C** | **Summary** | **Adjusted P Value** |
| --- | --- | --- |
| veh vs. veh + αIgD | ** | 0.0023 |
| Wortmannin (40nM) vs. wortmannin (40nM) + αIgD | ** | 0.0011 |
| Wortmannin (200nM) vs. wortmannin (200nM) + αIgD | ** | 0.0052 |
| veh vs. wortmannin (40nM) | ns | 0.49 |
| veh vs. wortmannin (200nM) | ns | 0.9675 |
| veh + αIgD vs. wortmannin (40nM) + αIgD | ns | 0.8761 |
| veh + αIgD vs. wortmannin (200nM) + αIgD | ns | 0.1153 |

**Supplementary Table 2:** Antibody-fluorochrome conjugates and other reagents used for flow cytometry in mouse experiments.

| Antibody target or reagent designation | Clone | Company | Conjugate | Dilution |
| --- | --- | --- | --- | --- |
| B220 | RA3-6B2 | BD Biosciences | V500 | 1/100 |
|  |  | Invitrogen | Qdot 655 | 1/100-400<br>(varies by lot) |
|  |  |  | APC | 1/200 |
| CD38 | 90 | eBioscience | Alexa Fluor 700 | 1/100 |
|  |  | BD Biosciences | Brilliant Violet 786 | 1/200 |
| CD45.1 | A20 | BD Biosciences | Alexa Fluor 647 | 1/200 |
|  |  |  | Alexa Fluor 700 | 1/100 |
| CD45.2 | 104 | BioLegend | Biotin | 1/400 |
|  |  |  | FITC | 1/100 |
|  |  |  | Brilliant Violet 785 | 1/100 |
| CD138 | 281-2 | BD Biosciences | Brilliant Violet 711 | 1/175 |
| Fixable Viability Dye eFluor 780 | N/A | eBioscience | N/A | 1/600-1000 |
| IgD | 11-26c.2a | BioLegend | PerCP-Cy5.5 | 1/200 |
|  |  |  | Alexa Fluor 700 | 1/100 |
| IgE | RME-1 | BioLegend | Unconjugated | 1/15 |
|  |  |  | FITC | 1/300-500 |
|  |  |  | PE | 1/300-500 |
| IgG1 | A85-1 | BD Biosciences | V450 | 1/300-400 |
|  |  |  | FITC | 1/300-400 |
| Ig$\lambda_{1,2,3}$ | R26-46 | BD Biosciences | FITC | 1/200 |
| IgM | II/41 | eBioscience | PE-Cy7 | 1/175-1/400 |
| NP | N/A | Conjugated in-house as described.<sup>10</sup> | APC | 1/750 of 0.1 mg/mL stock |
| phosphoS6 | N7-548 | BD Biosciences | Alexa Fluor 647 | 1/5 |
| PNA (peanut agglutinin) | N/A | Vector Laboratories | Biotin | 1/1000 |
|  |  |  | FITC | 1/500 |
| Streptavidin | N/A | Invitrogen | QDot 605 | 1/400 |
|  |  | BD Biosciences | Brilliant Violet 711 | 1/400 |
| TruStain FcX (anti-mouse CD16/32) | 93 | BioLegend | Unconjugated | 1/50 |

**Supplementary Table 3:** Antibody-fluorochrome conjugates and other reagents used for flow cytometry in human experiments.

| **Antibody target or reagent designation** | **Clone** | **Company** | **Conjugate** | **Dilution** |
| --- | --- | --- | --- | --- |
| CD20 | 2H7 | BioLegend | Pacific Blue | 1/200 |
| CD27 | O323 | BioLegend | PE | 1/50 |
| CD38 | HB-7 | BioLegend | PE-Cy7 | 1/200 |
| Fixable Viability Dye eFluor 780 | N/A | eBioscience | N/A | 1/600-1000 |
| IgD | IA6-2 | BioLegend | PerCP-Cy5.5 | 1/50 |
| IgE | MHE-18 | BioLegend | Unconjugated | 1/10 |
|  |  |  | APC | 1/150-300 |
| IgG1 | IS11-12E4.23.30 | Miltenyi Biotec | biotin | 1/150-300 |
| IgG4 | HP6025 | SouthernBiotech | FITC | 1/800 |
| IgM | MHM-88 | BioLegend | Brilliant Violet 605 | 1/125-1/200 |
| Streptavidin | N/A | Invitrogen | QDot 605 | 1/400 |
|  |  | BD Biosciences | Brilliant Violet 711 | 1/400 |
| Mouse gamma globulin | N/A | Jackson ImmunoResearch | Unconjugated | 1/100 |

## Acknowledgements

We thank J. Taunton and G.A. Smith for generously providing ibrutinib for our studies. We thank J.G. Cyster, K.M. Ansel, J. Zikherman, and J.F.E. Koenig for advice and comments on the study. This research was supported by the National Institute of Allergy and Infectious Diseases of the National Institutes of Health under Award Numbers R01AI130470 and R21AI154335; the Sandler Asthma Basic Research Center; and the Cardiovascular Research Institute at UCSF. C.D.C. Allen was a Pew Scholar in the Biomedical Sciences, supported by The Pew Charitable Trusts. A.K. Wade-Vallance was supported by a Doctoral Foreign Study Award from the Canadian Institute of Health Research, funding reference number DFD-170769. The content is solely the responsibility of the authors and does not necessarily represent the official views of the funding agencies.

